# Artificial neural network for brain-machine interface consistently produces more naturalistic finger movements than linear methods

**DOI:** 10.1101/2024.03.01.583000

**Authors:** Hisham Temmar, Matthew S. Willsey, Joseph T. Costello, Matthew J. Mender, Luis H. Cubillos, Jordan LW Lam, Dylan M Wallace, Madison M. Kelberman, Parag G. Patil, Cynthia A. Chestek

## Abstract

Brain-machine interfaces (BMI) aim to restore function to persons living with spinal cord injuries by ‘decoding’ neural signals into behavior. Recently, nonlinear BMI decoders have outperformed previous state-of-the-art linear decoders, but few studies have investigated what specific improvements these nonlinear approaches provide. In this study, we compare how temporally convolved feedforward neural networks (tcFNNs) and linear approaches predict individuated finger movements in open and closed-loop settings. We show that nonlinear decoders generate more naturalistic movements, producing distributions of velocities 85.3% closer to true hand control than linear decoders. Addressing concerns that neural networks may come to inconsistent solutions, we find that regularization techniques improve the consistency of tcFNN convergence by 194.6%, along with improving average performance, and training speed. Finally, we show that tcFNN can leverage training data from multiple task variations to improve generalization. The results of this study show that nonlinear methods produce more naturalistic movements and show potential for generalizing over less constrained tasks.

**Teaser:** A neural network decoder produces consistent naturalistic movements and shows potential for real-world generalization through task variations.

## Introduction

When asked about priorities for recovery using various neurotechnologies, people living with tetraplegia have consistently held restoration of upper-limb fine motor skills at the top (*1–3*). Over the past few decades, researchers have pursued this goal through the development of brain-machine interfaces (BMIs), which record neural activity through microelectrodes implanted in the brain and use a ‘decoder’ to map this activity to a control signal for prostheses. Using BMIs, human study participants have successfully produced text and speech (*4–6*), controlled computer cursors (*7*, *8*) robotic arms (*9*, *10*), prosthetic fingers (*11*), and, through muscle stimulation, controlled an individual’s own arm (*12*, *13*).

For the better part of the last two decades, linear decoders have dominated the state of the art in the BMI field (*14–19*). These approaches were motivated by studies showing apparent linear relationships between single-neuron firing rates and behavioral variables like kinematics of a reaching task or muscle activity of a precision grip (*20–24*). However, more recent work has suggested that the true relationship is more complicated: neural activity is driven at a population level by latent ‘neural trajectories’ through lower dimensional neural subspaces/manifolds, and these trajectories may have a highly nonlinear relationship to behavior (*25–31*). While linear decoders may be especially capable of predicting multi-DOF single-effector movements in constrained settings, recent studies have observed that linear decoders tested outside of their specific task may break down (*29*, *32*). As tasks become more sophisticated, the field had looked to nonlinear decoders as a potential solution for high-performance BMIs.

As a result, nonlinear models have now been used to significantly outperform linear methods in both offline and real-time BMI decoding tasks (*4*, *5*, *33–36*). In particular, nonlinear BMIs have recently enabled high-performance speech decoding in real-time (*6*, *37*). Recently, our group developed both a temporally-convolved feedforward neural network (**tcFNN**) which outperformed the ReFIT Kalman Filter and a recurrent neural network (RNN) which outperformed the tcFNN and a velocity Kalman Filter in a 2-degree-of-freedom (DOF) dexterous finger movement task (*38*, *39*).

While the use of artificial neural network (ANN) decoders has significantly improved BMI performance in tightly controlled experimental settings, it is unclear how well they will generalize to real-world use cases. As neural activity has been shown to shift significantly between variations to a task, it’s possible that ANNs may be sacrificing robustness to achieve such high-performance (*32*). A recent study demonstrated a recurrent neural network being effectively used to control two cursors simultaneously in real time, but it also demonstrated that without proper training, models may overfit to training datasets and fail during closed-loop control (*36*). Additionally, stochasticity (a.k.a. randomness) in ANN training approaches may lead to model instability, or variability in training, even when trained on the same data (*39*). Although an increasing number of studies have focused on adapting known ML architectures to neural decoding, there is a need to better understand whether ANNs will generalize and consistently converge to high-performing decoders. In this study, we use tcFNN as our ANN of choice, as the simplicity and feedforward architecture make it particularly amenable to examining precisely how these nonlinearities impact performance. In particular, we used tcFNN to predict the finger movements of a non-human primate in both open (offline) and closed-loop (online) experiments.

In this study, we show that offline, two nonlinear decoders (including tcFNN) outperform a linear decoder by producing a range of velocities that more closely matches those of natural hand finger movements. In addition, during real-time experiments, tcFNN maintains these improvements over the previous state-of-the-art linear approach (the ReFIT Kalman Filter). We then find that regularization techniques developed within the last decade enable tcFNN models to converge on high-performance solutions quickly and consistently across training runs, both offline and online. We also find that any remaining variance in tcFNN convergence is negligible during real-time control. Finally, we demonstrate that both linear and tcFNN decoders overfit to training data and struggle to generalize to task variations (‘contexts’). However, by including data from multiple contexts, this overfitting can be mitigated, and decoders can achieve near-indistinguishable performance from decoders trained and tested on the same context. We believe that these findings will allow researchers to improve the performance of their ANN decoders, regardless of their core architecture (e.g. RNN, CNN, transformer), as well as improving their generalization for real-world use.

## Results

### Improvements to the ‘fit’ of offline predictions using nonlinear decoders

To investigate how nonlinear decoders outperform linear approaches, we trained examples of both on four days of offline random target acquisitions from Monkey N. We used ridge regression (RR) as the representative linear decoder and two nonlinear decoders: a temporally-convolved feedforward neural network (tcFNN) developed by our group in 2022 and a dual-state movement/posture decoder developed by Sachs et al. in 2016 (*34*, *38*). Within each day, 5 models per decoder type were trained and then tested offline on a holdout set, for a total of n=20 decoders tested for each tcFNN, DS, and RR (n=60 decoders total). Example traces of index finger velocity predictions for each decoder are shown in Fig. 2A, with plots separated by decoder. We measured the average mean-squared error (MSE) of all models of each type across all four days (Fig. 2B), and found that mean tcFNN MSE was 28.0% lower (p < .01, paired t-test) than RR MSE, confirming our previous observations (*38*). DS also appears to produce lower error predictions, but mean DS MSE was only 5.7% lower (p < .01, paired t-test) than RR across all models and all days, still outperforming RR but to a lesser degree than tcFNN.

**Fig. 1.**
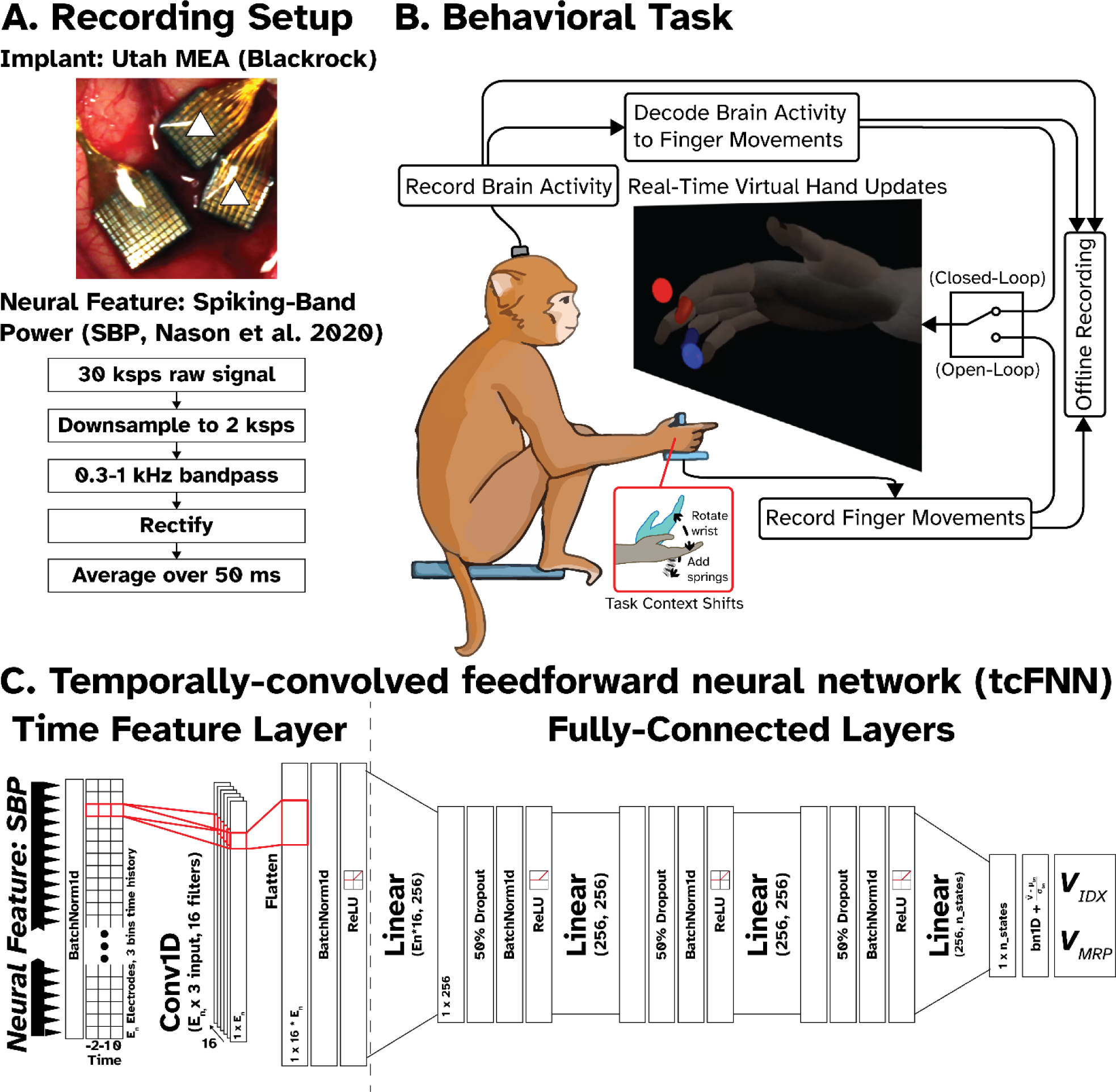
Experimental setup for finger movement BMI experiments. **(A)** *Top:* Utah arrays implanted in Monkey N. The two arrays marked with a white triangle are 8 x 8 Utah arrays connected to the same pedestal in the motor cortex (precentral gyrus). Recordings from these implants were used for decoding. Bottom: Flowchart describing neural feature extraction. 96 channels of 30 ksps raw data are down-sampled, filtered, rectified, and averaged, resulting in a low-power signal containing spiking information, called Spiking-Band Power (SBP, Nason et al. 2020) (*63*). **(B)** During an experiment, the monkey must move a virtual hand to match targets presented as spheres, 1 for each degree-of-freedom (DOF). During hand control (open-loop) trials, this is accomplished by moving the fingers in the manipulandum to match the same posture. The manipulandum measures finger flexion using flex sensors. In closed-loop (online) trials, the virtual hand is controlled by the output of the neural decoder, which maps brain activity to finger velocities. Additionally, for some trials, we introduced two ‘context shifts’ to the task, where either torsion springs were added to the flexion joint of the manipulandum to increase flexion force or the wrist angle was changed. **(C)** The primary decoder used in this study is a **t**emporally **c**onvolved **F**eedforward **N**eural **N**etwork (**tcFNN**), whose architecture is outlined above.

**Fig. 2.**
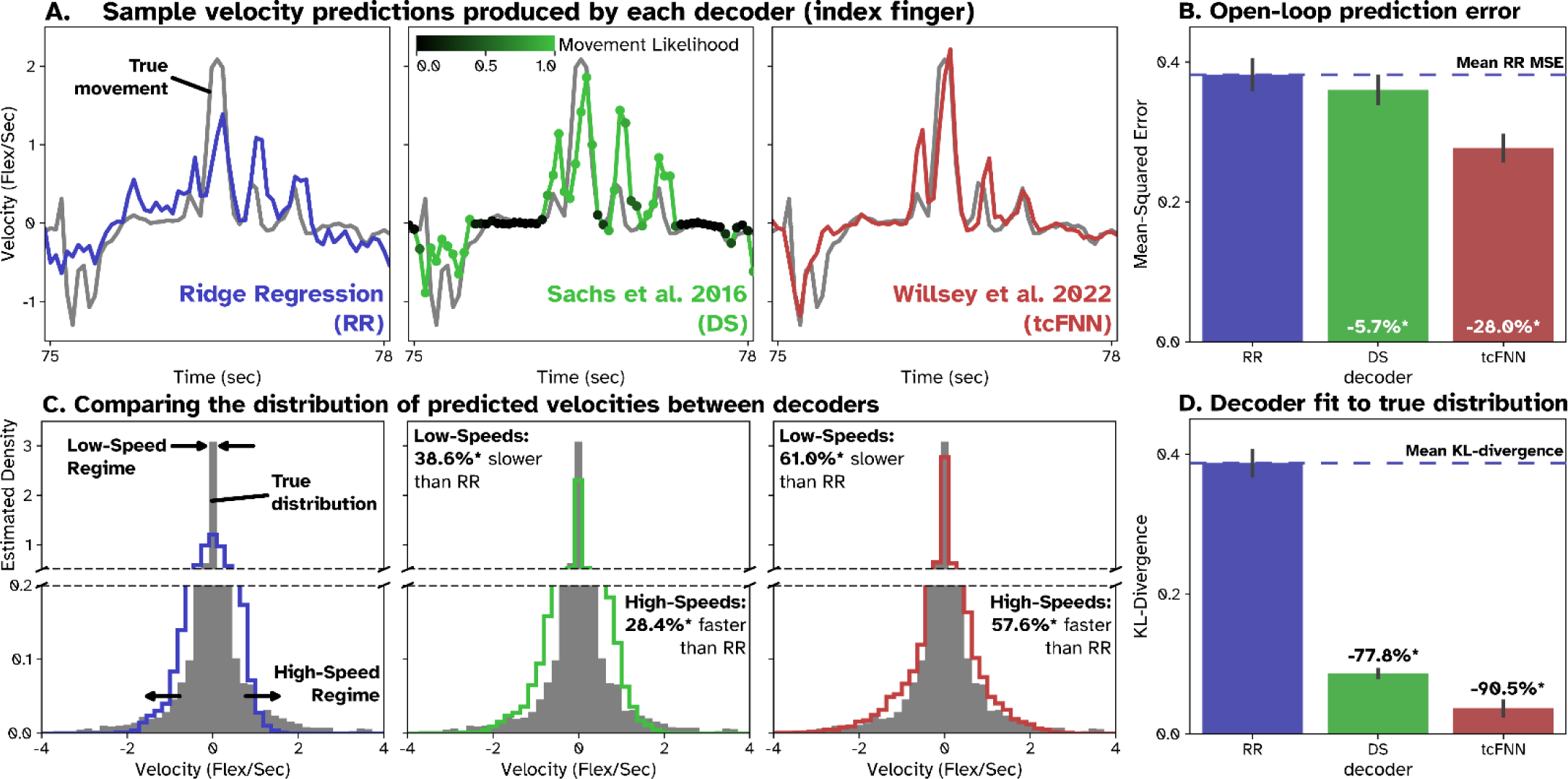
Distribution of nonlinear decoder predictions better matches open-loop movements. The three decoders compared here are Ridge Regression with 3 time bins of history (RR, blue), a dual-state movement-posture decoder proposed by Sachs et al. in 2016 (34) (DS, green), and tcFNN (red). True hand control (HC) is also shown in gray in subfigures A and C. **(A)** Sample predictions from each decoder on the same movement. **(B)** Average MSE of open-loop decoder predictions across four days on a 100-trial holdout set within each day. Gray lines show the standard error about each mean MSE. Asterisks (*) above a bar indicates p < .01 when comparing average MSE to RR, with the average % difference from RR across all decoders and days reported as well. **(C)** Example histograms of the distribution of velocities produced by actual hand movements in gray and by each decoder in their respective colors. Broken axes are used to highlight estimated density in high and low speed regimes. The percentage differences in the center/right panels are calculated by taking the percentage difference in mean speed at high and low speed regimes between RR and DS/tcFNN and then averaging across days. The high and low regimes are determined by selecting the top and bottom 10^th^-percentile of recorded speeds within a day, as well as the synchronized neural data for predictions. **(D)** Kullback-Leibler divergence (Kl-div) was calculated between scaled velocity distributions of hand control and decoder predictions, then averaged across folds and days, providing a measure of the fit or ‘distance’ between the two.

More specifically, both tcFNN and DS improved over RR in two clear ways: reaching higher speeds during peak movement and maintaining lower speeds during holding periods. To visualize these observations over time, we produced histograms of the velocity predictions for each model within each day. Single example histograms for each decoder type (all velocities from the test set on one day) are shown in Fig. 2C, color-coded by decoder. The filled gray histograms show the true velocities produced by hand control. We noted that the tcFNN histogram has larger tails than the RR histogram, indicating tcFNN produced a wider range of velocities than RR. The tcFNN histogram also shows a higher, tighter peak than RR, indicating tcFNN produced a higher proportion of low velocities than RR. To quantify these differences, we compared mean speed and MSE of tcFNN predictions to RR in the top and bottom 10^th^ percentiles of recorded movements, referred to as the high-and low-speed regimes of movement. In the high-speed regime, mean tcFNN predictions across all days and models were 57.6% faster (p < 0.01) and had 35.8% lower mean MSE (p < .01) than RR. TcFNN also improved RR in the low-speed regime, with 61.0% slower (p < 0.01) mean velocity predictions and 69.4% lower MSE (p < 0.01). Together, these results confirm that tcFNN moves substantially faster and slower than RR. Visually, the DS histogram also improves over the RR histogram in similar ways to tcFNN, but to a lesser degree. In the high-speed regime, mean DS predictions across all models and days were only 28.4% faster (p < 0.01) and had 16.5% (p < 0.01) lower MSE than RR. In the low-speed regime, mean DS predictions across all models and days were 38.6% slower (p < .01) and had 17.1% (p < .01) lower MSE than RR.

Next, we investigated which ‘distribution’ of velocities produced by either tcFNN or RR more closely resembled the true distribution of velocities during hand control. To do so, we calculated the KL-divergence between scaled HC histograms and decoder histograms on each day, as shown in Fig. 2D. Across days, we found that the mean KL-divergence from the true distribution was 90.5% lower (p < 0.01) for tcFNN than RR. DS histograms similarly showed improved KL-divergence from HC compared to RR, averaging 77.8% lower (p < .01) KL-divergence across models and days for DS predictions than RR predictions. Overall, these results suggest that tcFNN more closely fits the ‘true distribution’ of recorded velocities than RR, producing faster and slower predictions when it needs to, and generating velocity predictions which more closely resemble naturalistic control. It also significantly outperforms the DS decoder overall. This suggests that the presence of nonlinearities allows for more naturalistic and improved predictions offline than linear methods.

### ReFIT-tcFNN improves fit to true distribution during real-time control

While we have previously demonstrated that Re-tcFNN outperforms the linear RK online (*38*), it is unclear if this is due to the same improvements observed offline in Figure 2. To find out, we looked at three days of real-time comparisons between Re-tcFNN and RK with Monkey N. Example position traces for hand control (HC), RK, and Re-tcFNN are shown in Fig. 3A, with the IDX position in yellow and the MRS position in purple. During experiments, we observed that Re-tcFNN appeared to better resemble the high acceleration ‘snappy’ movements produced by true hand control than RK, suggesting Re-tcFNN may produce more naturalistic movements. To confirm this, we produced velocity histograms across all RK (blue), Re-tcFNN (red), and HC (gray) within each day (histograms for one day are shown in Fig. 3B). Across days, the distribution of Re-tcFNN velocities more closely resembled true movements than RK, with a 53.1% lower average KL-divergence from hand control (HC) than RK. Per-day KL-divergence can be seen in Table 2, along with trial counts. This result suggests that Re-tcFNN produces more naturalistic movements than RR, like what we observed offline between tcFNN and RR, although with a smaller relative difference between Re-tcFNN and RK divergences than tcFNN and RR divergences offline.

**Fig. 3.**
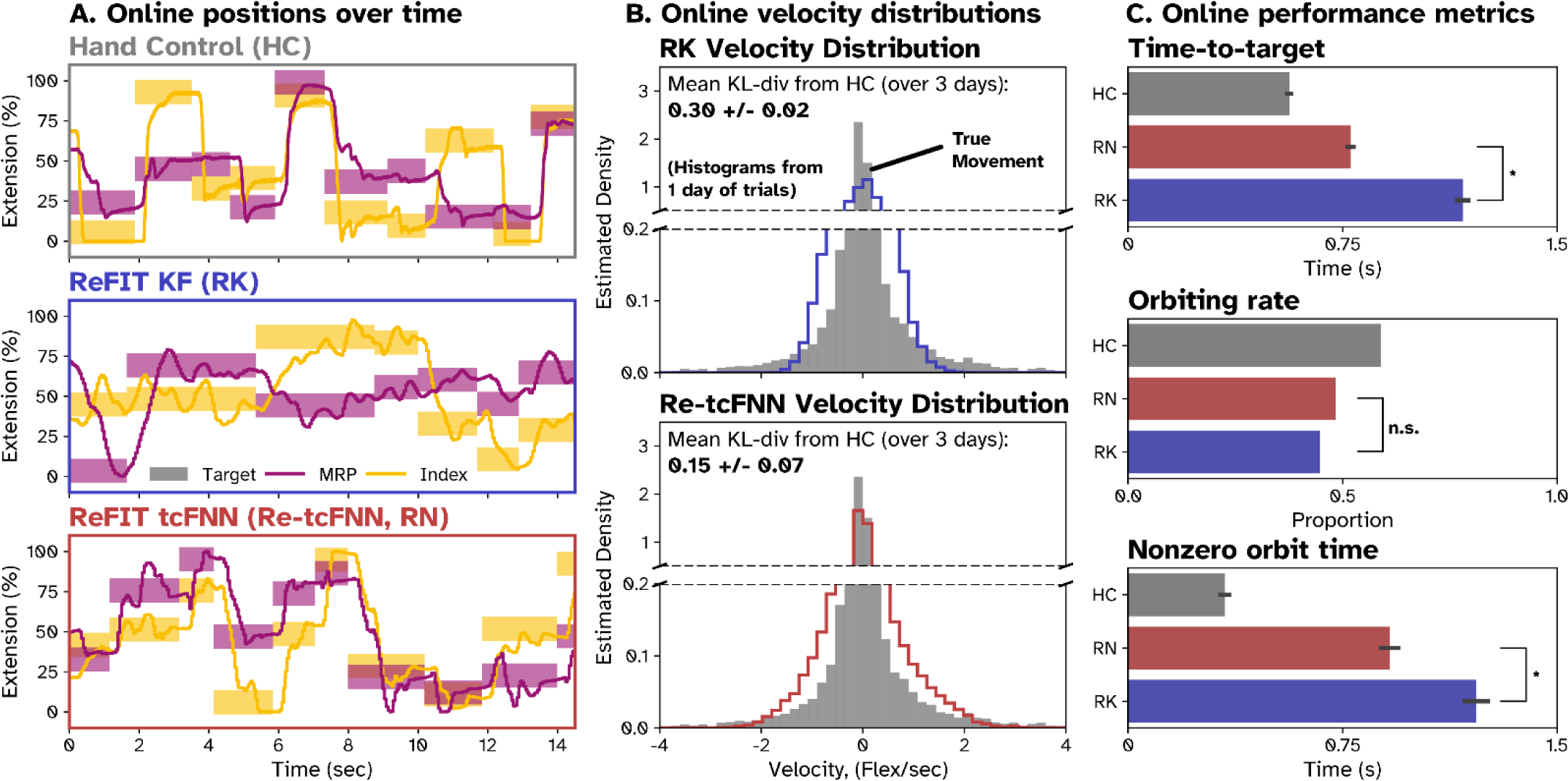
Comparing decoder fit to hand control movements in real-time control. **(A)** Position of the virtual hand across sample trials during HC, RK control, and Re-tcFNN control. Finger groups are differentiated by color (IDX yellow, MRS purple). Units are in % extension, covering the full movement range. **(B)** Example histograms showing the distribution of velocities produced by actual hand movements and real-time predictions on one day. The gray histograms show a representative distribution of true velocities using all hand-controlled training trials for the day. The blue and red histograms show the distributions of all RK and Re-tcFNN trials for the day, respectively. Mean and SD of KL-divergence from the HC histogram across days for each decoder is also shown. **(C)** Per-trial online performance metrics evaluating the abilities of each decoder to reach and stop in the target, averaged across all trials and all days. From top to bottom: Mean time-to-target (excluding trials with zero TT), orbiting rates, and nonzero orbiting time for HC (gray), Re-tcFNN (red), and RK (blue). Error bars denote standard error (across trials and days). Asterisks (*) indicate Re-tcFNN has significantly lower (p < .01, one-tailed two-sample t-test) average time-to-target and nonzero orbiting time than RK. N.S. indicates no significant difference found in a one-tailed difference of proportions test for orbiting rate between RK and Re-tcFNN across all trials on all days.

**Table 1.**
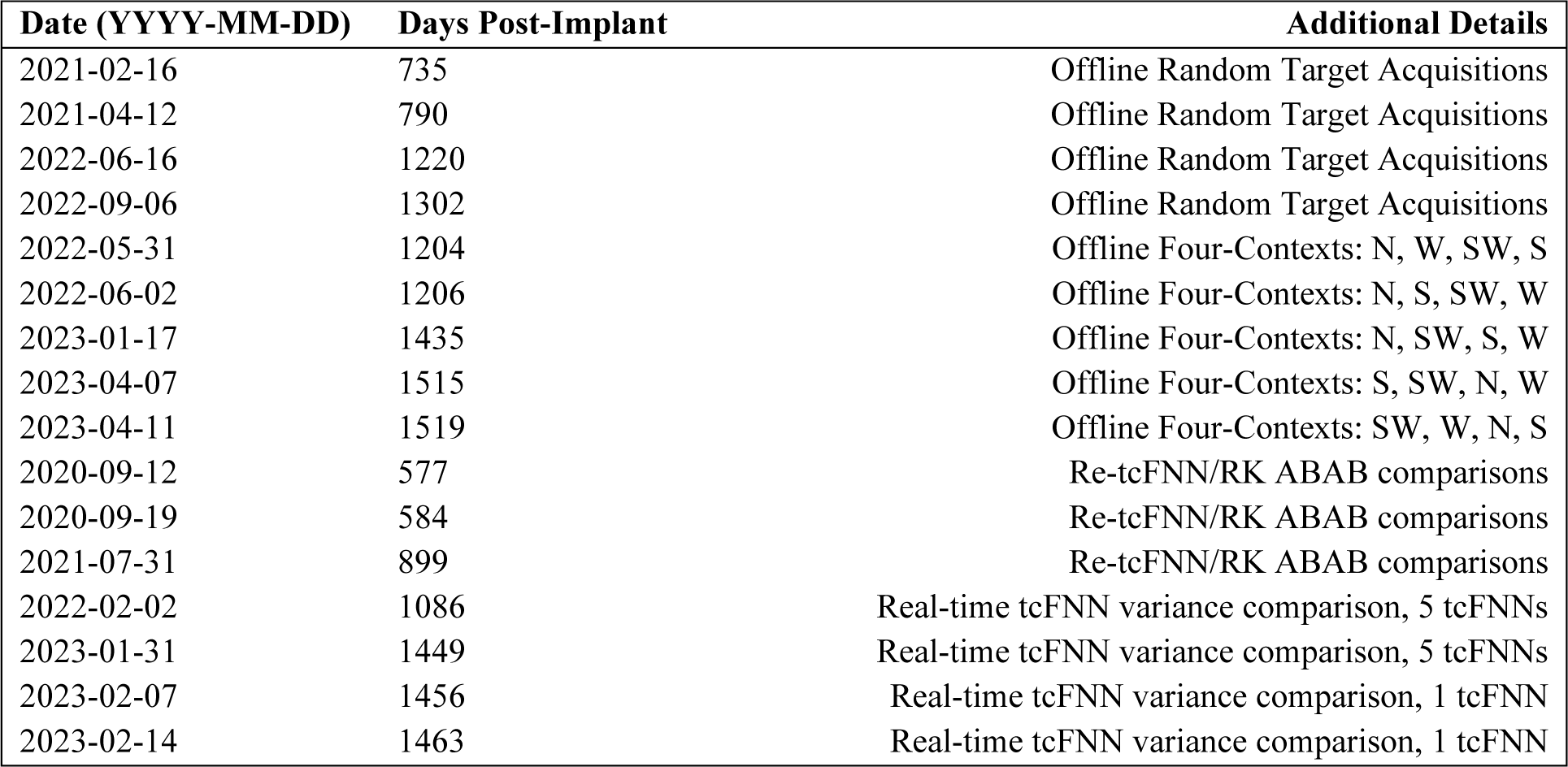
List of datasets. Each row indicates an experimental session used in the study. Columns show the date of the session, the number of days post implant, a short description indicating what kind of experiment was performed. These are further described in the Datasets section of the Methods and Materials. In the details for the Offline Four-Context sessions, single letter abbreviations indicate the order in which the blocks of context trials were performed (N = Normal, W = Wrist, SW = Spring-Wrist, S = Spring)

| Date (YYYY-MM-DD) | Days Post-Implant | Additional Details |
| --- | --- | --- |
| 2021-02-16 | 735 | Offline Random Target Acquisitions |
| 2021-04-12 | 790 | Offline Random Target Acquisitions |
| 2022-06-16 | 1220 | Offline Random Target Acquisitions |
| 2022-09-06 | 1302 | Offline Random Target Acquisitions |
| 2022-05-31 | 1204 | Offline Four-Contexts: N, W, SW, S |
| 2022-06-02 | 1206 | Offline Four-Contexts: N, S, SW, W |
| 2023-01-17 | 1435 | Offline Four-Contexts: N, SW, S, W |
| 2023-04-07 | 1515 | Offline Four-Contexts: S, SW, N, W |
| 2023-04-11 | 1519 | Offline Four-Contexts: SW, W, N, S |
| 2020-09-12 | 577 | Re-tcFNN/RK ABAB comparisons |
| 2020-09-19 | 584 | Re-tcFNN/RK ABAB comparisons |
| 2021-07-31 | 899 | Re-tcFNN/RK ABAB comparisons |
| 2022-02-02 | 1086 | Real-time tcFNN variance comparison, 5 tcFNNs |
| 2023-01-31 | 1449 | Real-time tcFNN variance comparison, 5 tcFNNs |
| 2023-02-07 | 1456 | Real-time tcFNN variance comparison, 1 tcFNN |
| 2023-02-14 | 1463 | Real-time tcFNN variance comparison, 1 tcFNN |

**Table 2.** Real-time performance comparison between Re-tcFNN and RK on three days. Re-tcFNN is abbreviated to RN in the table. Mean Time-to-target and nonzero orbiting time are included along with the respective SD and number of trials per decoder within each day. Orbiting rate (OR) and KL-divergence from hand control are also reported with the total number of trials included for each decoder on each day. Since overall KL-divergence is measured by averaging the three measurements on other days, # trials is not included.

| Performance Metrics: |  | Time-to-Target, ms |  |  | Nonzero Orbit Time, ms |  |  | Orbiting Rate |  | KL-Div from HC |  |
| --- | --- | --- | --- | --- | --- | --- | --- | --- | --- | --- | --- |
|  |  | Mean | SD | # trials | Mean | SD | # trials | Rate | # trials | Div | # trials |
| Day 1 | RK | 1355 | 737 | 373 | 1563 | 1107 | 193 | 0.51 | 382 | 0.29 | 382 |
|  | RN | 739 | 399 | 437 | 1173 | 935 | 270 | 0.60 | 451 | 0.16 | 451 |
| Day 2 | RK | 1298 | 702 | 362 | 1212 | 835 | 133 | 0.36 | 372 | 0.32 | 372 |
|  | RN | 843 | 391 | 476 | 787 | 512 | 192 | 0.39 | 489 | 0.21 | 489 |
| Day 3 | RK | 864 | 330 | 375 | 857 | 581 | 183 | 0.48 | 385 | 0.29 | 385 |
|  | RN | 742 | 317 | 381 | 664 | 458 | 181 | 0.47 | 388 | 0.07 | 388 |
| Overall | RK | 1171 | 654 | 1294 | 1217 | 926 | 509 | 0.45 | 1139 | 0.30 | N/A |
|  | RN | 778 | 377 | 1110 | 914 | 744 | 643 | 0.48 | 1328 | 0.15 | N/A |

From the examples in Fig. 3A, we observed that the more ‘naturalistic’ control of Re-tcFNN provided Monkey N with two distinct advantages over RK: First, Monkey N was able to reach targets more quickly with Re-tcFNN than RK. which can be seen in the examples shown in Fig. 3A, as the slopes of the position traces when moving from one set of targets to another are steeper in Re-tcFNN than RK. To quantify this, we measured mean time-to-target for all HC, Re-tcFNN and RK trials across all three days, as shown in Fig. 3C. Re-tcFNN had 31.6% faster mean time-to-target than RK (one-tailed two-sample t-test, p < .01) across days. Mean time-to-target, standard deviations, and trial counts for each day can be seen in Table 2. This lower time-to-target suggests Re-tcFNN reached targets more quickly than RK, outperforming RK in the high-speed regime.

The second advantage of Re-tcFNN was that it corrected overshoots more quickly than RK, although Re-tcFNN appeared to overshoot more often than RK. This can also be seen in Fig. 3A, as we see more oscillation around the targets before the final acquisition in the RK examples than the RN examples. To quantify Monkey N’s ability to stop in targets once reached, we measured both the proportion of trials that showed orbiting (orbiting rate, OR) and the average orbiting time of those trials (nonzero orbiting time, OT) for all HC, Re-tcFNN, and RK trials on all 3 days. Per-day OR and OT are listed in Table 2 (including trial counts), and averages across days are shown in Fig. 3C. A difference of proportions test across all trials on all days found no significant difference (p > .01) in orbiting rate between Re-tcFNN and RK, although both had lower orbiting rates than HC. Across days, Re-tcFNN averaged 27.5% lower nonzero orbiting time than RK (p < .01, two-tailed, two-sample t-test). Together, these results suggest that while Re-tcFNN and RK orbit targets at similar rates, Re-tcFNN allows Monkey N to correct overshoots/orbiting more quickly and settle into the target faster than RK, thus maintaining lower speeds than RK, similar to the differences observed between RR and tcFNN in Figure 2.

### Modern regularization techniques improve consistency of tcFNN convergence

One concern with tcFNN and ANNs in general is the stochastic methods used to train them, which could lead to models trained on identical training data converging on inconsistent solutions. To ensure that tcFNN training is consistent, on four days of offline random target acquisitions, we trained n=100 models of tcFNN on identical training data and tested on a hold-out dataset, seen in Fig. 4A. While there is some variance, no model deviates significantly from the rest, nor from the true movement (black line). We hypothesized that modern ANN training strategies were responsible for this consistency, including batch normalization (batchnorm) and neuron dropout. To this end, we also trained n=100 models each of three ‘ablated’ variations to the tcFNN model on the same training data for each day, removing batch normalization (noBN), neuron dropout (noDP) and both (noReg), shown in Fig. 4A. To compare overall performance between ablated models and tcFNN, we compared MSE across all days (Fig. 4B). Relative to tcFNN, the noBN and noReg models 88.2% and 69.3% higher had average MSEs (one-tailed, p < .01) than the tcFNN model, respectively. The noDP model had a 0.21% lower MSE (one-tailed, n.s.). During training, we observed that noDP and tcFNN models achieved higher performance in 10 epochs than the noBN and noReg models achieve in 80 epochs, seen in Fig. 4C. These results suggest that batchnorm contributes heavily to the improved performance and training speed of the tcFNN.

**Fig. 4.**
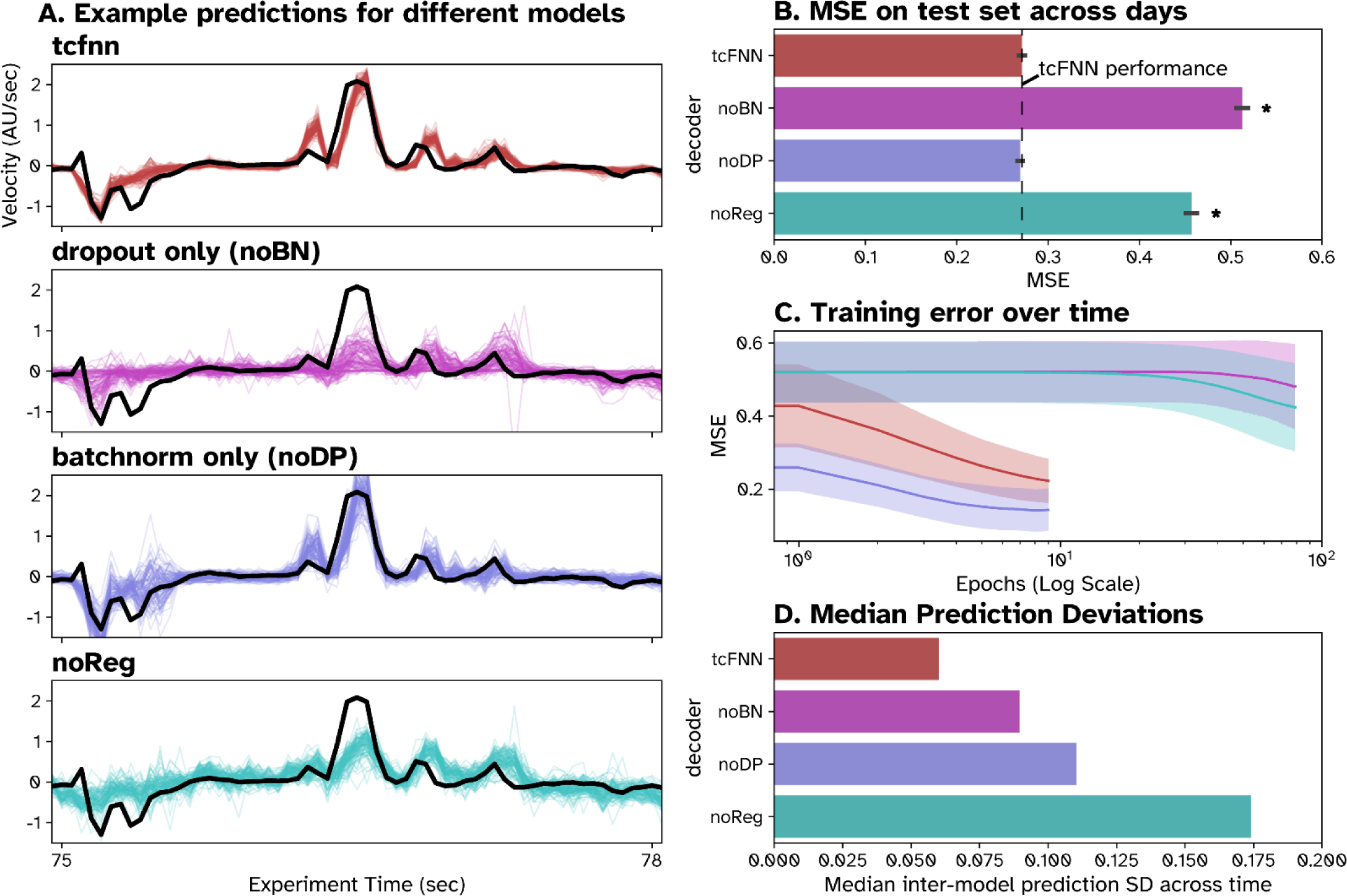
Contribution of different ML regularization techniques to tcFNN performance. **(A)** Example traces of 100 tcFNN (red), noBN (pink), noDP (blue) and noReg (turquoise) models trained on identical data compared to true hand velocities (black). **(B)** Bar plot showing mean MSE across all models of each type over all days, with standard error about the mean as black bars. Asterisk (*) indicates significantly (p < .01) higher average MSE than tcFNN. **(C)** Semilog plot of MSE on the training set at each training epoch for each model type. The solid line shows the average training error across all instances on all days for each type, while the shaded area shows the standard deviation. **(C)** Average prediction deviation across all predictions on all days for each decoder type. Median prediction deviation was measured by first taking the standard deviation across all models of one type at each timepoint in the test prediction, then taking the median across all time points on all days.

In Fig. 4D, we quantified the observed ‘spread’ of predictions for each model type using the median prediction deviation to assess how consistently each model type converged on similar solutions. Across days, tcFNN had the lowest deviation amongst its models, suggesting it has the most consistent performance. The noBN, noDP, and noReg models had 46.7%, 87.9% and 194.6% higher median deviation across days than tcFNN respectively. This suggests that batchnorm and neuron dropout contribute to model consistency, but neuron dropout contributes the most, and that the noReg model is extremely inconsistent in its convergence. Overall, we found that batchnorm and neuron dropout allow tcFNN to train quickly, reach high performance, and consistently converge on the same solution.

Since batch normalization contributed heavily to speeding up tcFNN training time, we hypothesized that by simply normalizing both the neural and behavioral data in our training set, we might see similar improvements. Thus, we trained 100 examples of each model with normalized inputs and normalized outputs for 15 epochs (Fig. S1A). In Fig. S1B, we see that normalization greatly reduced average MSE on the test data for noBN and noReg models (which had no previous normalization) while the tcFNN and noDP performances only changed slightly. Training times also improved for noBN and noReg, reaching similar training errors as tcFNN and noReg (respectively) within 20 epochs (Fig. S1C). Additionally, normalization improved the median prediction deviations of the noBN and noReg models, while median deviation increased slightly for tcFNN models and greatly for noDP models (Fig. S1D). While noReg and noBN had the lowest average MSEs and median deviations, respectively, tcFNN had lower average MSE than noBN and lower median prediction deviation than noReg. This highlights a tradeoff between training consistency and higher average performance which the tcFNN’s regularization strikes a balance between. Additionally, these results suggest that input and output normalization serve a similar purpose as batch normalization, improving overall performance and training times, and contributed more to reducing variance in decoder convergence than batch normalization.

### Online stability of tcFNN

Observing that offline consistency is improved by using regularization techniques, we next aimed to evaluate performance consistency during online, closed-loop trials. To assess this, on two days, Monkey N performed successive blocks of trials using five tcFNNs (without applying ReFIT) trained on identical training data. Additionally, we performed a control on two separate days, where Monkey N performed five consecutive blocks of trials using only one tcFNN. Throughput (bit rate) was used as a measure of trial performance. Each plot in Figure 5 shows box plots summarizing the per-trial throughput of each trial block within each day. Since stochastic initialization of each tcFNN instance could contribute to variance in decoder convergence, ReFIT recalibration was not applied, so overall performance is lower than observed in previous studies(*38*). To test for any major differences in performance between trial blocks in and across days, we performed a nested ANOVA, a hierarchical two-way ANOVA with each session/day as the high-level factor and intra-day trial block as the low-level factor. The nested ANOVA found a significant effect of day (p < .01) on average bitrate but found no significant effect (p > .01) of intra-day trial block nor a significant interaction (p < .01) between day and intra-day trial block. This means that average overall performance changed significantly from day-to-day, which we have observed previously, but that within each day, on average, there was no significant difference between trial blocks. Additionally, the lack of a significant interaction between the two factors suggests that there is no significant effect of each specific day on the average between trial blocks. If the days in which different models trained on the same data introduced significantly more variability, we would expect to see some significance either in the intra-day trial block factor or in the interaction. Thus, we concluded that the variance in tcFNN convergence does not introduce significant changes in model performance.

**Fig. 5.**
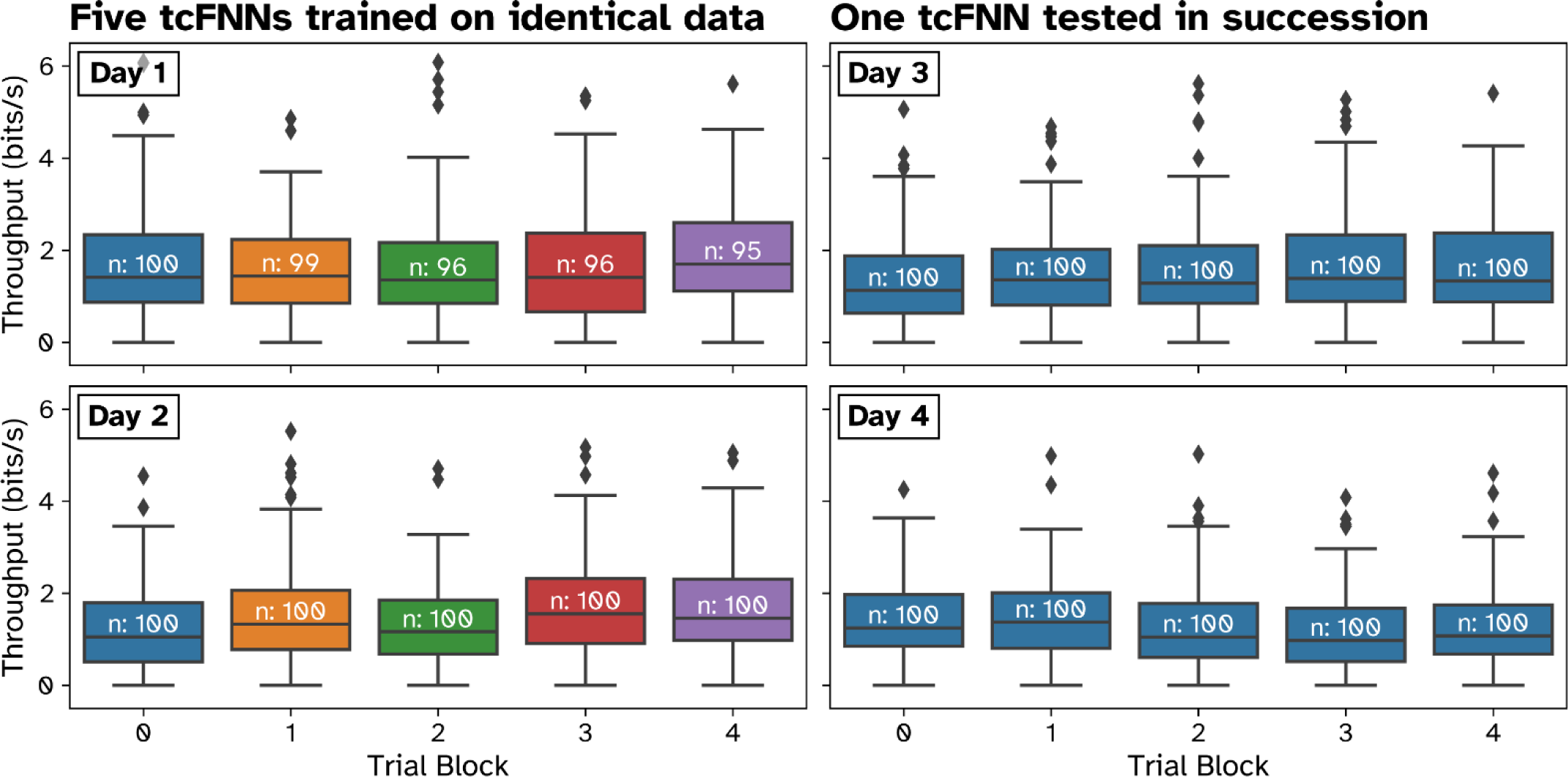
Validating stability of tcFNN (non-ReFIT) decoders trained on identical training datasets in real-time control. Color indicates a different decoder tested (within each day) **Left Column:** On 2 days (1 per plot), five tcFNN decoders were trained on identical training data (within a day) and tested in succession. Per-trial throughput was measured in bits/s. Each set of box plots shows the throughput median, quartiles, and IQR for each decoder tested (indicated by color) within a day. **Right Column:** On two separate days, a single tcFNN was tested six times in succession as a control experiment, simulating decoder switches after ∼100 trials.

### Impact of task changes to offline tcFNN predictions

It is possible that the high performance observed by current BMI decoders is effectively due to decoders overfitting to the highly controlled conditions imposed by research environments, which could render current approaches useless in real-world environments. To evaluate overfitting to changes in task context, we collected 600+ trials with Monkey N of four contexts on five days– normal, spring, wrist, and spring-wrist, and split them into training and testing datasets (as described in the Methods). Within each day, we trained tcFNN and RR models on each training set (including mixed datasets with data from multiple contexts) and tested them on four offline single-context testing datasets never seen during training. Fig. 6A shows the mean MSEs of RR and tcFNN decoders trained and tested on different combinations of contexts on one day. Compared across each training/testing combination on all days (5 predictions per combination, for a total of 600 predictions each across all days for RR and tcFNN), RR showed 21.2% higher MSE on average (p < .01) than tcFNN, suggesting that tcFNN outperforms RR across all combinations. We also observed that both decoders performed best when tested on the same context as they were trained on (*on-*context), as indicated by the downward-pointing triangles in Fig. 6A. We grouped MSEs into ‘on-context’ and ‘off-context’ predictions across all days, shown in Fig. 6B, and found that average off-context MSE for RR and tcFNN was 11.7% and 20.3% higher (one-tailed t-test, p < .003 Bonferroni-adjusted) than on-context MSE respectively. This suggests that both RR and tcFNN overfit to the task context, but relatively speaking, tcFNN overfits more to the task than RR. However, average off-context tcFNN MSE was still 9.1% lower than average on-context RR MSE (one-tailed t-test, p < .003, Bonferroni-adjusted), suggesting that while tcFNN may exhibit relatively more overfitting to context, it still learns more about the general task than RR.

**Fig. 6.**
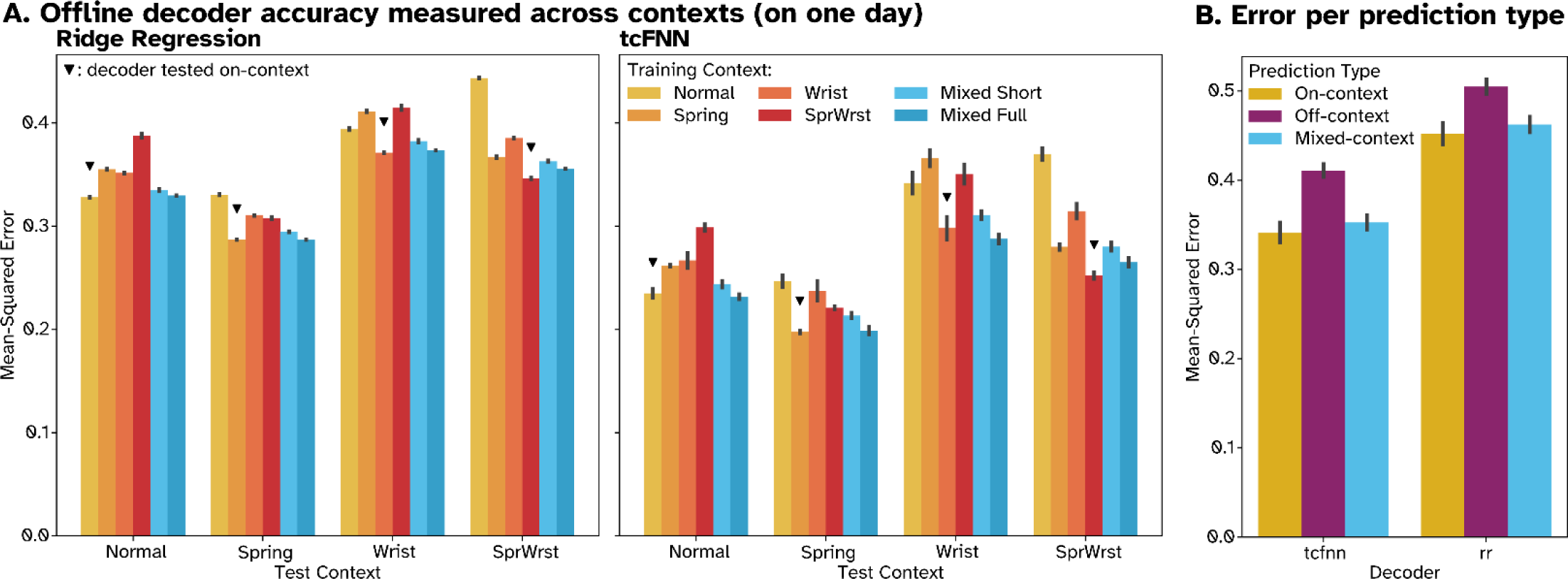
Measuring impact of context shifts on open-loop predictions for RR and tcFNN decoders across days. On five days, five ∼400 trial training datasets were prepared for Monkey N performing trials in each of four contexts: Normal, Spring, Wrist, and Spring-Wrist. Additionally, ten additional datasets were constructed per day, five ‘Mixed Short’ and five ‘Mixed Full’, each containing ∼100 trials and ∼400 trials from each of the four contexts, respectively. tcFNN and RR decoders were then trained on individual datasets and then tested on 100 held-out trials of each single context. **(A)** Average MSE for each combination of decoder training and testing context within one day, across five decoders per bar. Colors represent the source (training) context, and groups represent the target (test) context. In each group, a downward triangle indicates decoders trained and tested on the same context. Standard error about the mean is shown as black bars. **(B)** Average decoder MSE grouped into ‘on-context’, ‘off-context’, and ‘mixed-context’ predictions. On-context predictions (yellow) refer to predictions made on the same context as a decoder’s training data (downward triangle in A). Off-context predictions (purple) refer to predictions made on a different context as a decoder’s training data. Mixed-context (blue) predictions refer to predictions made by a decoder trained on the composite dataset.

Ideally, decoders could generalize to unseen contexts without modification, but if we relax this constraint slightly, we can consider using decoders which generalize after training on additional contexts. To this end, we trained several ‘mixed’ context decoders containing data from all 4 contexts, seen in Fig. 6A and grouped across days in Fig. 6B. Across all days, the mixed-context decoders outperformed the off-context decoders, with average MSEs 8.4% and 14.2% lower than off-context MSE for RR and tcFNN respectively (p < .003, Bonferroni-adjusted). In fact, the difference between mixed context MSEs was statistically insignificant from on-context performance, averaging only 2.3% and 3.3% higher than on-context MSE for RR and tcFNN respectively (p > .003, Bonferroni-adjusted). Overall, these results show that both decoders have improved potential for generalization when multiple contexts are included in the training data, and that tcFNN may benefit more from this strategy than RR.

Since training on multiple full-length datasets from each context is infeasible as the number of contexts increases, we created two types of mixed datasets, ‘short’ and ‘full’. The ‘short’ datasets contain ∼100 trials of data from each of the four contexts, resulting in a ∼400 trial dataset which is the same length as the single context datasets. The ‘full’ datasets contain ∼400 trials from each of the four contexts, resulting in a ∼1600 trial dataset four times as large as the single context datasets. Across all days, the average MSE of decoders trained on the ‘short’ datasets were actually 6.4% and 12.8% *lower* (for RR and tcFNN respectively) than average MSEs for decoders trained on ‘full’ datasets (p < .01, 100 short and full predictions within each decoder across days). This is counterintuitive to the idea that more data is always better, but regardless, this result shows that both RR and tcFNN may not require larger datasets but instead richer datasets to generalize across contexts, although real-time testing of these decoders is required.

## Discussion

Nonlinear decoders, and in particular artificial neural networks, are quickly gaining traction in movement and speech BCIs for their improved performance functionality and ‘generalizability’ (*4*, *8*, *35–37*, *40*). While these algorithms have been shown to improve over linear approaches, little is known about how these algorithms achieve their performance gains. We show that our tcFNN, likely representative of other nonlinear algorithms, decodes naturalistic, high-speed finger movements by closely matching the velocity distribution of actual finger movements. We also illustrate how recent regularization techniques are important to achieving reliable, high performance with artificial neural networks. Furthermore, these algorithms have the potential to generalize across a variety of context changes, a step in the right direction towards decoders robust to the constant context shifts imposed by everyday use. These findings provide insight into how to develop and to leverage neural networks for high-performance BCI applications.

This work suggests that ***nonlinear approaches to decoding produce more naturalistic control leading to higher performance BMIs.*** Multiple studies have shown that nonlinear BMI decoders can outperform linear methods in offline and online settings (*33–35*, *38*, *39*), and we showed that tcFNN and DS showed improvements over RR consistent with these results. More strikingly however, in Figure 2 we also showed that the flexibility of tcFNN allowed it to better imitate real finger movements, allowing it to produce a distribution of velocities closer to natural control than RR. This resulted in a decoder which substantially improved over RR in both high and low-speed regimes, without needing to sacrifice one for the other. While the simpler nonlinearity of the DS decoder also showed similar improvements over RR, it did not reach the same level of performance as tcFNN. However, these analyses were performed offline, and it’s been observed that high-accuracy offline decodes do not always translate to good real-time performance (*36*). Because of this, Figure 3 showed that even in real-time experiments, Re-tcFNN provides more naturalistic control than RK, the previous state-of-the-art linear approach. More naturalistic performance may encourage the use of nonlinear decoders in real-world BMI devices, particularly if improved functionality and ease-of-use lead to reduced device abandonment, as issues with these have been shown to contribute heavily to device abandonment for devices like upper-limb prosthetics (*41*).

We also demonstrate here that ***modern deep learning (DL) techniques allow neural networks to quickly and consistently converge on high-performance solutions.*** The stochastic optimization methods used during tcFNN training introduce potential for inconsistent decoder performance, even when trained on the same data. Fortunately, in the last decade, techniques like batch normalization, neuron dropout, Kaiming initialization, and Adam optimization have been shown to empirically improve performance and reduce overfitting and training times (*42–45*). In this study we showed that batch normalization allowed for improved performance and faster training times, and neuron dropout reduced the variance in converged solutions. Together, these two approaches allow for tcFNN to train quickly, effectively, and consistently. Furthermore, we showed that any remaining variance in tcFNN convergence does not impact online performance within a day. If numerically optimized nonlinear decoders are the future of BMI, it will be important to demonstrate that they converge consistently onto similar solutions. Further investigation using comparative techniques like projection-weighted canonical correlation analysis will help show if the internal structure of these networks converges as consistently as their outputs (*46*).

Next, we showed that ***DL decoders provide a potential avenue to tackle BMI generalization across variations to a task*.** Overfitting to training data limits BMI generalization across different conditions, as we observed in the decreased off-context performance of both tcFNN and RR decoders. It is known that DL decoders tend to overfit more strongly than analytical approaches, and we did observe relatively higher overfitting of tcFNN to the task context than RR. However, even an ‘overfit’ tcFNN outperformed RR at its best. Additionally, both decoders showed potential for improved generalization when trained on multiple contexts, and it’s likely that tcFNN will see further improvements as the number of contexts increases. This capability of the algorithm to learn across multiple contexts is likely similar to how artificial neural network decoders have been found to be robust to neural instabilities over time (*35*). Research into data augmentation strategies like noise injection and new training paradigms like continual and multi-task could improve tcFNN’s robustness and improve cross-context performance (*47–50*). Two recent studies have shown that incorporating transfer learning into decoder training improves stability over time and allows for cross-species decoding (*51*, *52*).

In this work, we explored ‘*how*’ nonlinear decoders improve over linear methods but did not address underlying reasons ‘*why*’. While some research suggests that individual neural firing rates are linearly correlated with behavior (*22*, *24*), many others also support a highly nonlinear relationship between behavior and neural activity in the motor cortex (*25–28*, *53*). Additionally, spinal interneurons in the motor pathway provide places for nonlinear integration of information end effector (*54*, *55*). Thus, it’s possible that feedforward neural networks may serve as a more biologically relevant decoder, but it’s also likely that they have no biological relevance and instead achieve greater performance due to their flexibility as universal approximators (*56*). A truly biologically relevant model would need to account for many feedforward and recurrent inputs from several brain areas (*57–60*).

In this study we showed that artificial neural network decoders show promise as high-performance naturalistic decoders for brain machine interfaces. However, there are multiple limitations to our study. Primarily, these limitations stem from having only a single NHP subject, performing only a single task involving two degrees of freedom. These findings may not translate to other NHP subjects, or with different and more complex tasks. Additionally, we only used two nonlinear decoders as representative of nonlinear decoders in general. We also did not perform any optimization on the tcFNN training hyperparameters or architectures, instead using the values used in our 2022 study by Willsey et al. (*38*). Future work may further explore our findings with different tasks, and in humans.

We have demonstrated that nonlinear decoders provide more naturalistic control over a motor brain machine interface than linear approaches. We also showed that neural network decoders using standard regularization strategies quickly converge on consistent, high-performance solution, and that with the proper training data, they may generalize across tasks. Thus, artificial neurla networks have the potential to revolutionize motor brain-machine interfaces for people living with spinal cord injury and greatly improve their quality of life.

## Materials and Methods

### Experimental Design

This study investigated the ability of nonlinear decoders to outperform current linear methods as well as to investigate the potential of artificial neural network decoders to converge consistently and generalize to task context. The experiments were designed to collect synchronous brain and behavioral data during a two degree-of-freedom (DOF) finger movement task, as well as to test the performance of BMI decoders on said task. This was a single non-human primate study, and no power analysis was calculated before the study. The protocols in this study were approved by the Institutional Animal Care and Use Committee at the University of Michigan.

### Implants

One non-human primate (NHP), Monkey N (NHP, Rhesus macaque, *Macaca mulatta*) was implanted with Utah microelectrode arrays (Blackrock Neurotech, Salt Lake City, UT) in the hand area of the precentral gyrus, as described previously (*16*, *32*, *61*, *62*). Monkey N was implanted with two 64-channel arrays in the right hemisphere of motor cortex (precentral gyrus), (seen in Fig. 1A, the two arrays marked with triangles). Simultaneous recording from multiple channels was limited to 96 channels. The quality of recordings from each channel was inspected visually every day during experiment setup. Channels with low activity or visible noise were excluded from real-time decoding, however in offline decoding analyses, all 96 channels were used. Recordings from Monkey N included in this study were captured between 578 and 1519 days post-cortical implant.

### Decoding Hardware and Neural Feature Extraction

Signals from the implanted Utah array were recorded in real time through the Cerebus Neural Signal Processor (NSP) (Blackrock Neurotech, Salt Lake City, UT). The brain-machine interface is an xPC Target (Mathworks, Natlick, MA) based system. In this study, we used spiking-band power (SBP) as our neural feature, where raw 30 ksps data is downsampled to 2 ksps, a 300-1000 Hz bandpass filter is applied, and the resulting value is rectified, as described previously (*63*, *64*). This 2 ksps SBP feature is then further averaged over 50-ms into non-overlapping bins for decoder training and testing. In real-time experiments, binned SBP data was sent to a separate Linux machine which decoded the neural data and sent back the predicted kinematics.

### Behavioral Task

Monkey N was trained to sit in a chair (Crist Instruments, Hagerstown, MA) and perform a dexterous finger movement task. During the task, the monkey was presented with a monitor showing a virtual hand, which was controlled either by their physical hand in a manipulandum or by using decoded neural activity (Fig. 1B). In hand-control mode (termed “HC”), moving the index (IDX) or middle-ring-small (MRS) finger groups controlled the corresponding virtual finger groups. Offline analyses refer to predicting the finger kinematics from the neural activity using HC. In brain-control mode (“online” control), neural activity was decoded to predict virtual hand movement in real-time. Hand movement data was captured using flex sensors placed in a manipulandum in which the monkey’s hands were placed (FS-L-0073-103-ST, Spectra Symbol, Salt Lake City, UT). The actual task performed by the monkey is to match pseudo-randomly presented targets for each finger group (IDX or MRS), as described in previous studies (*16*, *38*).

The monkey received a juice reward for successful trials. Hold times for successful trial completion were 750 ms in ‘training’ mode and 500 ms in ‘testing’ mode. The failure condition was a trial timeout of 10 s, except for days 1449, 1456, and 1463 post-implant, which was 5 s. Target size was 15% of the movement range. Positions for each degree-of-freedom (DOF) will be referred to either as %flexion or %extension, and velocity units are in flex/sec, where ‘flex’ is the proportion of the total movement range from 0 (full extension) to 1 (full flexion) for either degree of freedom. This means that positive velocities are flexion and negative velocities are extension. On some days we shifted the ‘context’ of the task either by adding springs to the manipulandum to increase the required flexion force or by rotating the wrist angle, both of which have previously been shown to modulate neural activity while maintaining similar kinematics (*32*). Applying these shifts results in four task ‘contexts’: the *spring* context, the *wrist* context*, spring-wrist* context (both simultaneously), and *normal* context (no shifts).

### Datasets

Below, we describe the datasets used in analyses throughout the study. Additionally, Table 1 reports the date, post-implant day, monkey, and a brief description of each session used in the study.

#### Offline random target acquisitions

On four days, Monkey N performed 1000+ trials of the random target finger movement task. Data for each day was split into five 400-trial training datasets and one 100 trial test dataset according to the leave-one-out approach described in Offline Decoding Analyses.

#### Offline four-context random targets

On five days, Monkey N performed 600+ trials of the random target task for each of the four contexts previously described to evaluate decoder performance across contexts. Data for each context (and within each day) was split into five 400 trial training datasets and one 100-trial testing dataset, according to the leave-one-out approach described in Offline Decoding Analyses. Additionally, we prepared two types of ‘mixed context’ training datasets, ‘full’ and ‘short’, as described in Offline Decoding Analyses, resulting in 10 mixed context datasets per day.

#### Real-time Re-tcFNN/RK ABAB comparisons

On three days first reported in Willsey et al. 2022, Refit tcFNN (Re-tcFNN) and the ReFIT Kalman Filter (RK) were compared in ABAB comparisons, as described previously (*38*).

#### Real-time tcFNN variance comparisons

Four total days of online control experiments were performed for this dataset, with the goal of evaluating the variance in performance of online decoders trained on identical data. On 2 days we trained 5 tcFNN decoders on a 400-trial random target dataset (within each day) and tested them in successive 100+ trial blocks. On 2 additional days, we similarly trained a single tcFNN decoder and tested it in five 100+ trial blocks. Since the stochastic initialization of tcFNN parameters could contribute to variance in decoder performance, we chose not to apply the ReFIT procedure for these real-time comparisons, therefore the decoders used on these days are tcFNN, not Re-tcFNN.

### Decoders

Throughout this study, we compare the offline predictions and real-time performance of multiple linear and nonlinear approaches to decoding. Each of these approaches is described in more detail below. Offline trials are also referred to as ‘hand-control’ (HC) trials. In this study, we use the words ‘model’ and ‘decoder’ interchangeably.

#### Ridge Regression (RR) (65)

RR was used as a representative example of an offline linear decoder. While similar to linear regression, ridge regression includes L2 regularization, which aims to balance model performance with the magnitude of model weights, as determined by a hyperparameter lambda. A lambda value of .001 was used.

#### ReFIT Kalman Filter (RK) (18)

Kalman filters makes use of a linear dynamical model to incorporate prior knowledge of estimated behavior and the current observation to predict the true value of a hidden state, such as the velocity of two finger groups controlled by a brain-machine interface. The ReFIT KF improves on the basic Kalman Filter by implementing a recalibration stage in real-time control to improve performance. The ReFIT Kalman Filter implementation used in this study is described in Nason et al. 2021 (*16*).

#### Dual State Decoder (DS) (34)

Sachs et al. 2016 proposed a dual-state decoding paradigm, which predicts movement velocities by taking a weighted linear combination of two linear regressions trained on either high-speed movements or low-speed movements (referred to as the ‘movement’ and ‘posture’ states in the original study). The weights are determined by a modified LDA classifier, which is trained to predict the likelihood of being in either the movement or posture state from the same neural data used in the linear regression. The classifier also features an adaptive threshold, which aims to maintain a pre-defined ratio by measuring the proportion of fast and slow classifications over a sliding window and adjusting accordingly. In this study, we chose a ratio of 1:1.

#### Temporally Convolved Feedforward Neural Network (tcFNN) (38)

The temporally convolved feedforward neural network (tcFNN), whose architecture is outlined in Fig. 1C, was first proposed by Willsey et al. in 2022. The same hyperparameters were used as in the previous publication. At the input, the network applies multiple time-wise convolutions across 3 time-steps of the neural feature to each channel independently, unlike a ‘traditional’ convolutional neural network (CNN), where the filter bank is iterated over parts of an entire signal (like passing over an entire image). The tcFNN was implemented using PyTorch with standard linear, batchnorm, and convolutional layers. Convolutional filters were trained on 3-bins of time history (150 ms total).

#### ReFIT Temporally Convolved Feedforward Neural Network (Re-tcFNN) (38)

The Re-tcFNN is a variation on the tcFNN which is trained using a similar recalibration stage as the RK, as fully described in Willsey et al. 2022. Briefly, after an initial online session, recorded online velocities pointed away from the target are ‘corrected’ to produce a ‘refit’ training data set. The original tcFNN is then fine-tuned on the ‘refit’ data for 500 iterations at a learning rate of 2e-4.

#### Non-regularized tcFNN

We also trained versions of tcFNN without batch normalization and artificial neuron dropout, two training strategies which have empirically been shown to help ANNs regularize (*42*, *43*). Otherwise, the same architecture, initialization, and optimization were used. These versions are called noReg (no dropout or batch normalization), noDP (batch normalization only), and noBN (dropout only). Together, these models are referred to as the ‘ablated’ tcFNN models. We did not perform additional hyperparameter optimization on the ablated models.

#### Additional notes on tcFNN architecture

All neural networks were trained in PyTorch to minimize MSE between predicted and true velocities using the Adam optimizer (learning rate 1e-4, weight decay 1e-2). While networks intended for closed-loop experiments were trained for 3500 iterations with triangular velocity redistribution as outlined in Willsey et al. 2022, offline networks were trained for 10 epochs with no velocity redistribution, with two exceptions. The noReg and noBN ablated models, which were trained for 80 epochs each, and all models trained on normalized neural and behavioral activity were trained for 15 epochs each. Additionally, tcFNN normalizes the final velocity output, which is then scaled back to true velocity ranges. In the 2022 study, this was performed by using the median velocity as an offset and scaling to training velocity peaks. In this study, besides the three days of Re-tcFNN/RK comparisons, normalized velocities are instead scaled by using the training data to train two independent ridge regressions (each with a single weight, single bias) to map from the normalized IDX/MRS predictions to the final IDX/MRS predictions.

#### Time history

For RR, DS, and all tcFNN variations, 100 ms (2 bins) of lookback (time history) were also included for each SBP feature. This means that each decoder was trained to predict velocities from 150 ms (3 bins) of averaged SBP on each channel.

In our offline analyses, we aimed to investigate the effect of nonlinearity on direct mappings from neural signals to behavior, so we opted to use tcFNN, DS, and RR as exemplar nonlinear (tcFNN, DS) and linear (RR) decoders. In real-time comparisons, direct mapping approaches like linear regression and tcFNN have been outperformed in closed-loop experiments by decoders like the ReFIT Kalman Filter (RK) and ReFIT-tcFNN (Re-tcFNN) (*18*, *38*, *66*). Because of this, we opted to use RK and Re-tcFNN as representative closed-loop linear and nonlinear decoders in the following comparison.

### Offline Decoding Analyses

#### Offline data preparation

On each day of offline trials, unsuccessful trials were removed, and the dataset was trimmed to the middle 600 trials. The final 100 trials in the 600-trial block were then held out and not seen by decoders until testing. The first 500-trial dataset was randomly shuffled and split into five equal length blocks. Then, using a leave-one-out method, we created five ∼400 trial training datasets (a.k.a. training folds) for each day. The remaining 100 trials for each fold were used as validation but were not included in testing datasets. On days where multiple contexts were tested, this procedure was followed within each context, producing five 400-trial training datasets and one 100-trial testing dataset per context. Furthermore, within each fold, we created two ‘mixed’ context datasets, ‘full’ and ‘short’. The full mixed datasets contain all the training data from each context within a fold, while the short mixed-context datasets contain only 25% of the data from each context. This resulted in five 1600-trial ‘full’ mixed datasets and five 400-trial ‘short’ mixed datasets.

#### Offline error

Offline prediction error was measured using the mean-squared error (MSE) between decoder predictions and true velocities. Since most analyses include predictions from multiple instances of similar decoders, MSEs are typically reported as the average MSE across all decoders of the same type and are accompanied with a standard error.

#### Comparing speed regimes

We compared the mean speeds and MSEs of offline decoder predictions in two ‘speed regimes’, high-and low-speed. Here, speed refers to the velocity magnitude within a single degree of freedom. We defined the high-speed and low-speed regimes within a day, as the time points containing top and bottom 10^th^ percentiles of true speeds (across both degrees of freedom), respectively. This allowed us to investigate both how decoders differ at high and low speeds and if these differences are closer to true hand movements.

#### Comparing distribution of predicted velocities

We measured the Kullback-Leibler divergence (KL-div) between the distribution true velocities and velocities generated by a decoder. PMFs were estimated by scaling histograms of velocities (across DOFs) to sum to one. When comparing the distributions of decoders tested in real-time, we used the histogram of the experimental day’s training data to calculate the histogram.

#### Median prediction deviation

We tested if the ablated (noBN, noDP, noReg) models converged less consistently on the solutions than tcFNN within a day. For each velocity measurement in the test dataset, we measured the standard deviation of velocity predictions across all instances of each model type. We took the median of the resulting Nx2 vector across all time points and degrees of freedom, resulting in a scalar measure of how much each model type varied in convergence per day. This was further averaged over days.

#### Context comparisons

On days with trials performed in different contexts, decoders were trained for each single-and mixed-context dataset. All decoders from one day were then tested on the four single-context testing datasets for that day. MSE of these predictions were then grouped by three prediction types, *on-context* (training context matches test context), *off-context* (training context does not match test context), and *mixed-context* (training contains multiple contexts).

### Online Decoding

#### Online data preparation

For all analysis of online trials, the first 5 trials of any contiguous block of online trials were excluded to allow Monkey N to become familiar with each decoder. Additionally, only the middle 100 trials of each block were included for analyses on the real-time tcFNN variance experiments, or all trials if there were <100 (lowest was 95). For analyses comparing RK to Re-tcFNN, all successful, closed-loop trials where the monkey was performing the task were included. In an initial real-time tcFNN variance experiment, we observed that Monkey N progressively improved over the course of a session. To reduce the influence of behavior factors, we had Monkey N perform an additional trial block at the end of each experiment using the first decoder tested, not included in analyses. For the single decoder variance experiments, this meant running 6 blocks of the same decoder. If there was a large difference in performance between trials performed in the first and last blocks of the session, we would exclude the day, however no days needed to be excluded except the initial experiment.

#### Performance metrics

General online performance was assessed using throughput as defined by Fitt’s law, as previously described (*38*), measured in bits per second. Time-to-target (TT) was used to assess a decoder’s ability to quickly reach the target, and was measured as the time from the start of a trial to the first time both finger groups entered their respective targets, as described previously (*16*). Orbiting time (OT) was used to assess a decoder’s ability to correct overshoots once targets were reached and was measured as the time from the first target acquisition (the endpoint of time-to-target) to the final time the targets are acquired by both finger groups before the hold time is completed, as described previously (*16*). Orbiting Rate (OR) was used to assess a decoder’s ability to prevent overshooting and was measured as the ratio of trials with nonzero OT to total trials performed by the decoder. In analyses investigating trial orbiting, both OR and non-zero OT are used.

### Statistical Analyses

In general, all statistical comparisons between decoder performances were performed by grouping all decoders of each type across all days. A two-sample one-tailed t-test was then performed, with an alpha level of 0.01. On the other hand, in the comparison between RR and tcFNN models trained and tested on different contexts, we performed a pairwise two-sample one-tailed t-test, matching each RR to the corresponding tcFNN trained on the exact same training and testing dataset. When comparing orbiting rates between Re-tcFNN and the ReFIT Kalman Filter, a difference of proportions test was used, as we are comparing proportions over a session (and across days) rather than multiple observations of trial-by-trial metrics. Additionally, to investigate the variance of multiple decoders trained on the same data online, we used a nested two-way ANOVA, with the two factors being the session number (date) and the run within the session (on the days with multiple decoders, each run is a different decoder).

## Acknowledgments

Thank you to Eric Kennedy for his animal and experimental support, Chris Andrews for his assistance with statistical analysis, the University of Michigan Unit for Laboratory Animal Medicine for their expert veterinary and surgical support, and the University of Michigan Biointerfaces Institute for their support.

## Funding

National Science Foundation grant 1926576 (HT, MJM, PGP, CAC, MMK)

National Science Foundation grant 2223822 (HT)

National Institutes of Health grant T32NS007222 (MSW, JLWL)

National Science Foundation GRFP grant 1841052 (JTC)

Agencia Nacional de Investigación y Desarrollo (ANID) of Chile (LHC)

Dan and Betty Kahn Foundation grant 2029755 (CAC, DMW)

Michigan Robotics Institute (DMW)

## Author contributions

Conceptualization: HT, MSW, JTC, CAC

Methodology: HT, MJM, MSW, JTC, LHC

Investigation: HT, JTC, MSW, MJM, DMW, MMK, LHC

Visualization: HT, LHC

Software: HT, JTC, MSW

Validation: LHC, JTC

Supervision: CAC, PGP

Writing—original draft: HT, JLWL

Writing—review & editing: HT, JTC, JLWL, MJM, MSW, DMW, MMK, PGP, CAC

## Data and materials availability

All data are available in the main text or the supplementary data. Along with the data presented here, datasets for all analyses will be made available for research purposes. Analysis code will also be made available through GitHub. Final materials will be released along with the publication of this manuscript.

