## Supplementary Figure 1 for "Artificial neural network for brain-machine interface consistently produces more naturalistic finger movements than linear methods"

Hisham Temmar et al.

### Supplementary Figures

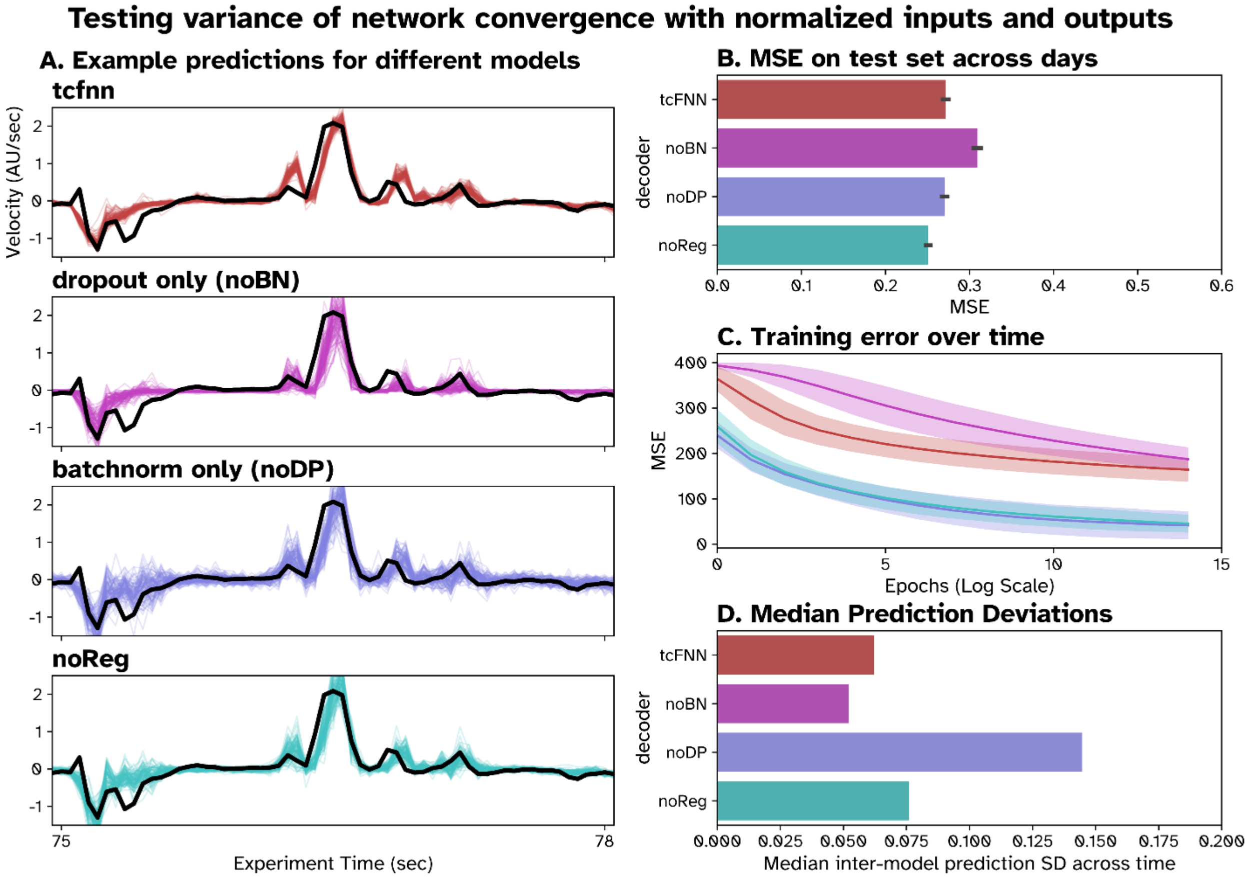


**Fig. S1.**

**Contribution of different ML regularization techniques to tcFNN performance with input and output normalization.** Here, tcFNN and three ablated models are trained on datasets with normalized inputs and outputs to evaluate the impact of data normalization on performance. **(A)** Example traces of 100 tcFNN (red), noBN (pink), noDP (blue) and noReg (turquoise) models trained on identical data compared to true hand velocities (black). **(B)** Bar plot showing mean MSE across all models of each type over all days, with standard error about the mean as black bars. **(C)** Plot of MSE on the training set at each training epoch for each model type. The solid line shows the average training error across all instances on all days for each type, while the shaded area shows the standard deviation. **(C)** Average prediction deviation across all predictions on all days for each decoder type. Median prediction deviation was measured by first taking the standard deviation across all models of one type at each timepoint in the test prediction, then taking the median across all time points on all days.
